# Cyclic constrained immunoreactive peptides from crucial *P. falciparum* proteins: potential implications in malaria diagnostics

**DOI:** 10.1101/2021.10.11.463910

**Authors:** Kapil Vashisht, Sukrit Srivastava, Vandana, Ram Das, Supriya Sharma, Nitin Bhardwaj, Anupkumar R Anvikar, Tong-Soo Kim, Byoung-Kuk Na, Ho-Joon Shin, Kailash C Pandey

## Abstract

Malaria is still a global challenge with significant morbidity and mortality, especially in the African, South-East Asian and Latin American region. Malaria diagnosis is a crucial pillar in the control and elimination efforts, often accomplished by administration of mass-scale Rapid diagnostic tests (RDTs). The inherent limitations of RDTs-failure of detection in low transmission settings, and deletion of one of the target proteins-Histidine rich protein (HRP) are evident from multiple reports; thus necessitating the need to explore novel diagnostic tools/targets. The present study used peptide microarray to screen potential epitopes from 13 antigenic proteins (CSP, <u>EXP1</u>, LSA1, TRAP, AARP, AMA1, <u>GLURP</u>, MSP1, <u>MSP2</u>, MSP3, MSP4, P48/45, HAP2) of *P. falciparum*. Three cyclic constrained immunoreactive peptides-C6 (EXP1), A8 (MSP2), B7 (GLURP) were identified from 5,458 cyclic constrained peptides (in duplicate) against *P. falciparum* infected sera. Peptides (C6, A8, B7-cyclic constrained) and (G11, DSQ, NQN-corresponding linear peptides) were fairly immunoreactive towards *P. falciparum*-infected sera in dot-blot assay. Using indirect ELISA, cyclic constrained peptides (C6 & B7) were found to be specific to *P. falciparum* infected sera and further, observed to be significantly reactive towards antibodies from field-collected *P. falciparum* infected sera. Notably, the structural location of the epitopes defines the reactivity, observed by the preferential recognition of cyclic constrained peptides *vs* linear peptides and corroborated by the homology modeling analysis of selected proteins. In conclusion, the study identified three cyclic constrained immunoreactive peptides (C6, B7 & A8) from *P. falciparum* secretory/surface proteins and two of them (C6 & B7) were validated for their diagnostic potential with field-collected *P. falciparum* infected sera samples.

## Introduction

The WHO malaria report 2020 reported an estimated ∼229 million malaria cases in 2019 compared to ∼228 million malaria cases in 2018. The slowing rate of decline in malaria cases since 2015 is worrisome, globally ^1^. Prompt malaria diagnosis is a key factor in the malaria control programs & elimination strategies, worldwide. In India, malaria diagnosis relies primarily on rapid diagnostic tests (RDTs), which are based on the target proteins-histidine rich protein (HRP) & lactate dehydrogenase (LDH) recognition, by specific antibodies in the infected sera ^2,3^. However, the reports of genetic deletions of *P. falciparum* (*Pf*hrp2/3) genes, are of great concern in malaria diagnosis and burden estimation efforts ^4^. Such evolutionary adaptation of the parasite to evade the most successful diagnostic modality is a crucial challenge. Further, the advent of drug resistance towards artemisinin (ART), the active ingredient of artemisinin combination therapies (ACTs), is also becoming a barrier for the country’s efforts on malaria control and elimination ^5^. Thus, it is imperative to explore novel diagnostic targets with high sensitivity and specificity for prompt malaria diagnosis.

The conserved secretory/surface proteins of the parasite proteome play crucial roles in the pathogenesis of malaria ^6–11^. Several full-lenth proteins such as circumsporozoite protein (CSP), P48/45, P25, P28, apical membrane antigen-1 (AMA1), glutamate rich protein (GLURP), exported protein-1 (EXP1), merozoite surface protein-1/2/3 (MSP1/2/3), schizont egress antigen (SEA1) etc. have been studied extensively for their immunogenicity and thus have been investigated as potential diagnostic targets ^12–14^. It is noteworthy that compared to whole proteins, immunogenic peptides inherently are cost-effective in synthesis, stability and handling; also offer possibility of chimeric synthesis from different proteins. A number of B-cell epitopes have been identified by microarray studies using proteins *viz*. SEA1, MSP7, AMA1, reticulocyte binding proteins (RBPs) from *P. falciparum* ^13^. The peptide microarray technology provides a good platform to screen potential immunogenic epitopes from multiple proteins. In the present study overlapping, cyclic constrained short peptides from multiple antigenic proteins of the *P. falciparum*, were screened. The cyclic constrained peptides would mimick 3D structure of the epitopes in the secondary structure of the proteins. Immunoreactive peptides from *P. falciparum* infected sera were identified and further validated using dot blot assays and ELISA techniques.

## Methods

### Screening of immunoreactive peptide epitopes

#### Design of the microarray slide

The candidate proteins (n=13) were selected from three major stages of parasite life-cycle-pre-erythrocytic, erythrocytic and sexual stages **(Table 1)**. The full-length amino acid sequences of the candidate proteins were mapped as short peptides (7 & 13 amino acids), as these lengths of peptides generally bind to Class-I and Class-II HLA during antigen presentation and developing immune response. The mapped peptides were arranged in an overlapping fashion of 4 & 10 amino acids, on the microarray slide. A total of 10,916 peptides (5,458 peptides in duplets) from 13 proteins were immobilized on the microarray slide in duplets. The immobilized peptides were cyclic constrained to mimic 3D conformation of the epitopes. Thioether bridging between -C and –N terminal amino acids was used to generate cyclic constraints. The polio control peptide tags (KEVPALTAVETGAT) were included in the microarray slide as positive controls. The microarray slide with immobilized peptides, mapped and designed as explained above, was commercially manufactured by PEPperPRINT^®^, Hamburg, Germany.

**Table 1:** List of selected proteins based on their antigenic nature and important role in *P. falciparum* pathogenesis. The selected proteins are expressed in various life cycle stages of the malaria parasite *P. falciparum*.

| S. No. | Name of proteins | Life cycle stage of *P. falciparum* |
| --- | --- | --- |
| 1. | CSP- circumsporozoite protein | Pre-erythrocytic (sporozoite) |
| 2. | EXP1- exported protein 1 |  |
| 3. | LSA1- liver stage antigen |  |
| 4. | TRAP- thrombospondin-related adhesion protein |  |
| 5. | GLURP- glutamate rich protein | Blood stage antigens (merozoite) |
| 6. | AARP- apical asparagine rich protein |  |
| 7. | AMA1- apical membrane antigen |  |
| 8. | MSP1- merozoite surface protein 1 |  |
| 9. | MSP2- merozoite surface protein 2 |  |
| 10. | MSP3- merozoite surface protein 3 |  |
| 11. | MSP4- merozoite surface protein 4 |  |
| 12. | P48/45 | Gamete surface (gametes) |
| 13. | HAP2 |  |

#### Sample collection

The *P. falciparum-*infected sera samples were collected after approval by the Institutional Ethics Committee (IEC), ICMR-National Institute of Malaria Research, Delhi, India [ECR/NIMR/EC/2019/308]. Whole blood samples (n=2) from *P. falciparum-*infected patients were collected from primary health center (PHC) Ujina, Mewat, Haryana, India. The samples were also confirmed by microscopy. Sera were separated from clotted blood by centrifuging the vacutainers at 1500 g for 5 minutes and stored at -20 °C until further use.

#### Antigenic peptides screening by peptide microarray immunoassay

The customized microarray slide with immobilized conformational peptides (5,458 peptides in duplets) as described in the previous section was commercially procured and assay was performed as per manufacturer’s instructions^15^. Briefly, the microarray slide was incubated for 15 minutes in PBS containing 0.05% Tween 20 (PBS-T, pH 7.4) and blocked with Rockland Blocking Buffer (RL) (Rockland Immunochemicals) at room temperature (RT) for 30 minutes. Further, the microarray slide containing pooled sera from *P. falciparum*-infected patients (1:100 dilution) in 10% RL/PBS-T, was incubated overnight at 4°C on an orbital shaker. The slide was washed thrice with PBS-T for 1 minute and further incubated with goat anti-human IgG (Fc)-Cy5 (Invitrogen) antibodies at 1:2500 dilution. Subsequently, the microarray slide was washed thrice with PBS-T for 1 minute, dipped in 1mM Tris pH 7.4 and dried with air stream. The microarray slide was scanned using microarray scanner (Molecular devices, USA) at Translational Health Sciences and Technology Institute (THSTI), Faridabad, Haryana, with a resolution of 20 μm and fluorescent read-out of Cy5 dye was recorded. The PepSlide analyzer software algorithm calculated median foreground intensities (background-deductedintensities) of the spots in duplicates.

### Validation of immunoreactive peptide epitopes

#### Dot blot immunoassay with cyclic constrained and linear peptides

To qualitatively analyze the reactivity profiles of selected peptides against patient serum antibodies, cyclic constrained and linear peptides were commercially synthesized. Purified recombinant *P. falciparum* GLURP protein (Glutamate rich protein) was used as a positive control. The *P. berghei* and *P. falciparum* lysates from *in-vitro* cultures were used to demonstrate *P. falciparum* specific immunoprecipitation of serum antibodies, and Casein was used as a negative control. Uninfected serum from a healthy volunteer was also used in the dot blot immunoassay. Briefly, 50 μg of peptides were coated onto PVDF membrane (0.45㲼m), using Bio-Rad dot-blot SF apparatus; post activation and equilibration with methanol and PBS, respectively. The peptide spots were dried using a vacuum pump and further incubated with *P. falciparum*-infected and uninfected sera samples (1:100 dilution), further blocked with 5% skimmed milk overnight at 4 °C. The strip was washed thrice with PBS and incubated with anti-human IgG secondary antibody (1:2500 dilution) for 1 hr. at room temperature. The blot strips were washed thrice with PBST (PBS + 0.05% Tween 20) and once with PBS alone. The blot was developed using DAB (3,3’-Diaminobenzidine tetrahydrochloride) substrate, where the development of brown precipitate was observed to identify the reactive peptides.

#### ELISA analysis of cyclic constrained epitopes

Direct-Enzyme-Linked Immunosorbent Assay (ELISA) was performed to validate and quantitatively analyze the reactive potential of the chosen peptides with antibodies in *P. falciparum* infected sera samples. Purified recombinant GLURP was used as a positive control, and BSA (Bovine serum albumin) was used as a negative control. Briefly, the 96-well ELISA plate was coated with 50 μg of peptides, purified recombinant GLURP and BSA diluted in 1X PBS and kept overnight at 4 °C. Afterwards, ELISA plate was aspirated and blocking buffer (5% casein + 0.1 N NaOH in 1X PBS) was added to each well followed by incubation for 2 hrs at 4 °C. Next, diluted (1:100) *P. falciparum* infected patient sera were added in each well and incubated at 4 °C for 2 hrs. After the primary antibody incubation from sera, the ELISA plate was washed with wash buffer (0.05% Tween-20 in 1XPBS). The ELISA plate was further incubated for 1 hr. with anti-human IgG secondary antibody tagged with HRP with a dilution of 1:3000 in 1X PBS buffer. Next, the ELISA plate was again washed thrice with wash buffer. The ELISA plate was further developed by 30 minutes incubation at room temperature with peroxidase substrate solution [*o*-Phenylenediamine dihydrochloride (OPD) + sodium citrate (pH 5.0) + H_2_O_2_]. The peroxidase reaction with OPD produced a dark yellow product indicating the titer of secondary antibodies bound to the serum (primary) antibodies. The ELISA plate was scanned and analyzed at 405 nm for quantitative analysis.

## Results

### Screening of immunoreactive peptide epitopes

#### Screening by peptide microarray immunoassay

A total of thirteen surface and secretory proteins from *P. falciparum* were chosen due to their antigenic potential **(Table 1)**. A schematic representation of the peptide microarray slide design for immunoreactive epitope mapping is shown in **Figure 1**. The microarray slide was scanned using a microarray scanner and images were superimposed on the grid for data read-out and analysis. Every fluorescent spot corresponded to the immobilized peptide of the microarray slide. The fluorescent intensities of the spots indicate the relative immunogenicity of the corresponding peptide, quantified as green foreground mean **(Figure 2)**. A heat map of the highest scoring peptides of the microarray slide, covering epitopes from all the proteins was generated **(Figure 3)**. Out of the 104 identified peptides, there were three highest scoring peptides from the antigenic proteins GLURP, EXP1 and MSP2. These three peptides were further selected for further validation studies **(Table 2)**.

**Figure 1:**
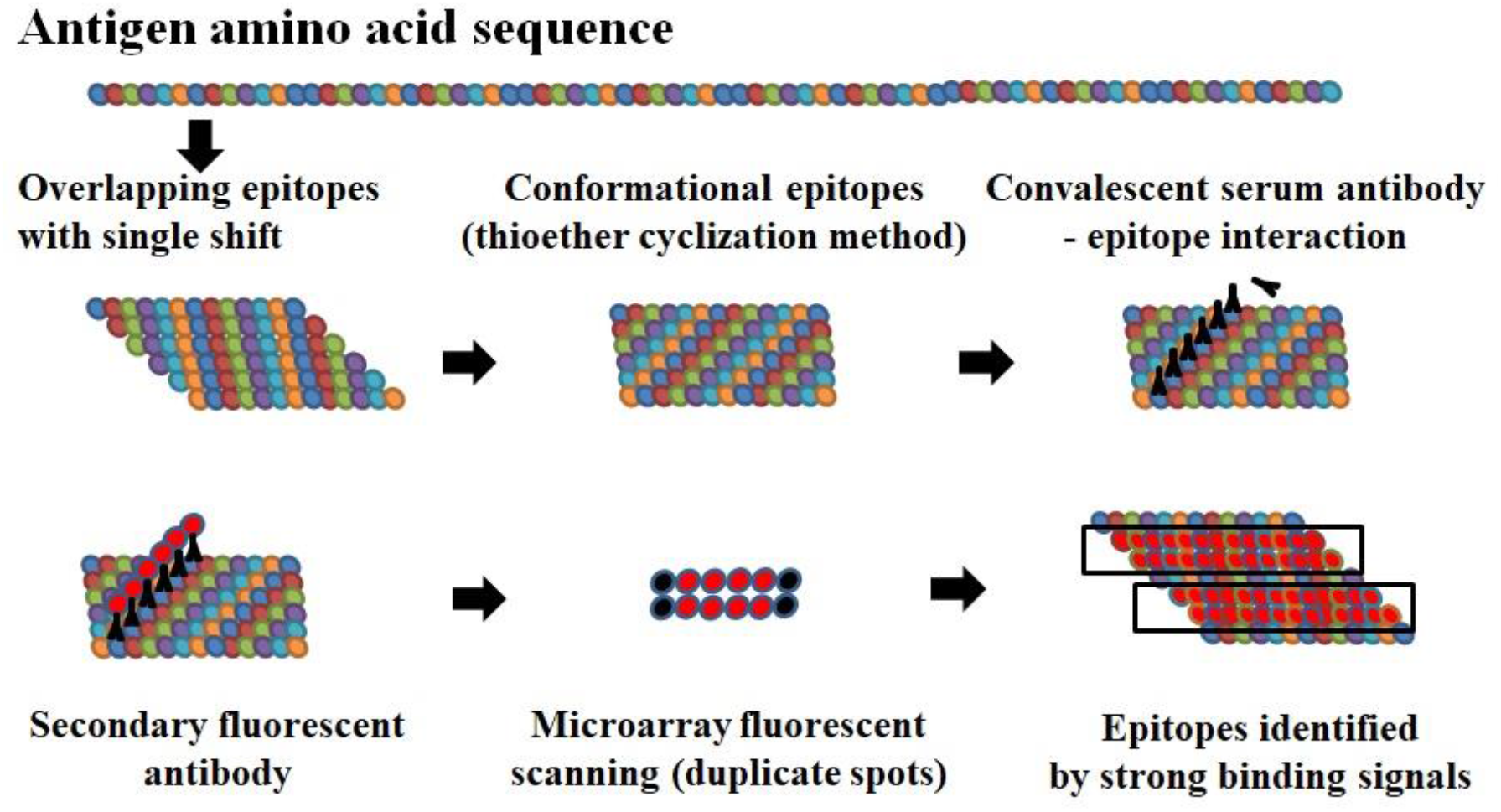
Schematic representation of the peptide microarray design with overlapping conformational peptides for antigenic peptide epitope mapping. The chain of circles shows peptides, indicating an overlapped arrangement of the short peptides from the antigenic proteins.

**Figure 2.**
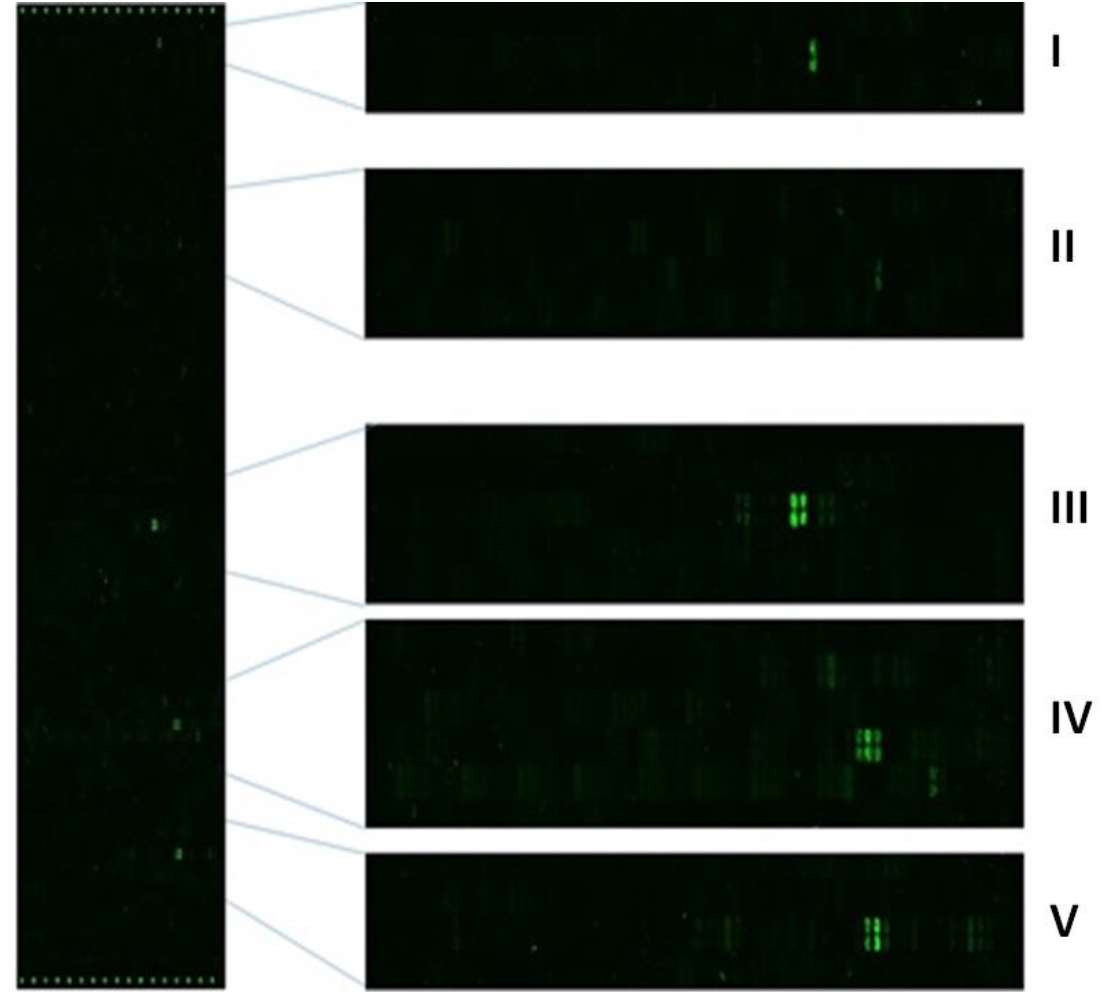
The fluorescent image of microarray epitope mapping showing multiple peptides interacting with the antibodies in patient sera. The zoomed cut-outs of fluorescent spots with reactive epitopes are shown in I to V lanes. Fluorescent spots on the top and bottom borders represented the polio positive control peptides.

**Figure 3:**
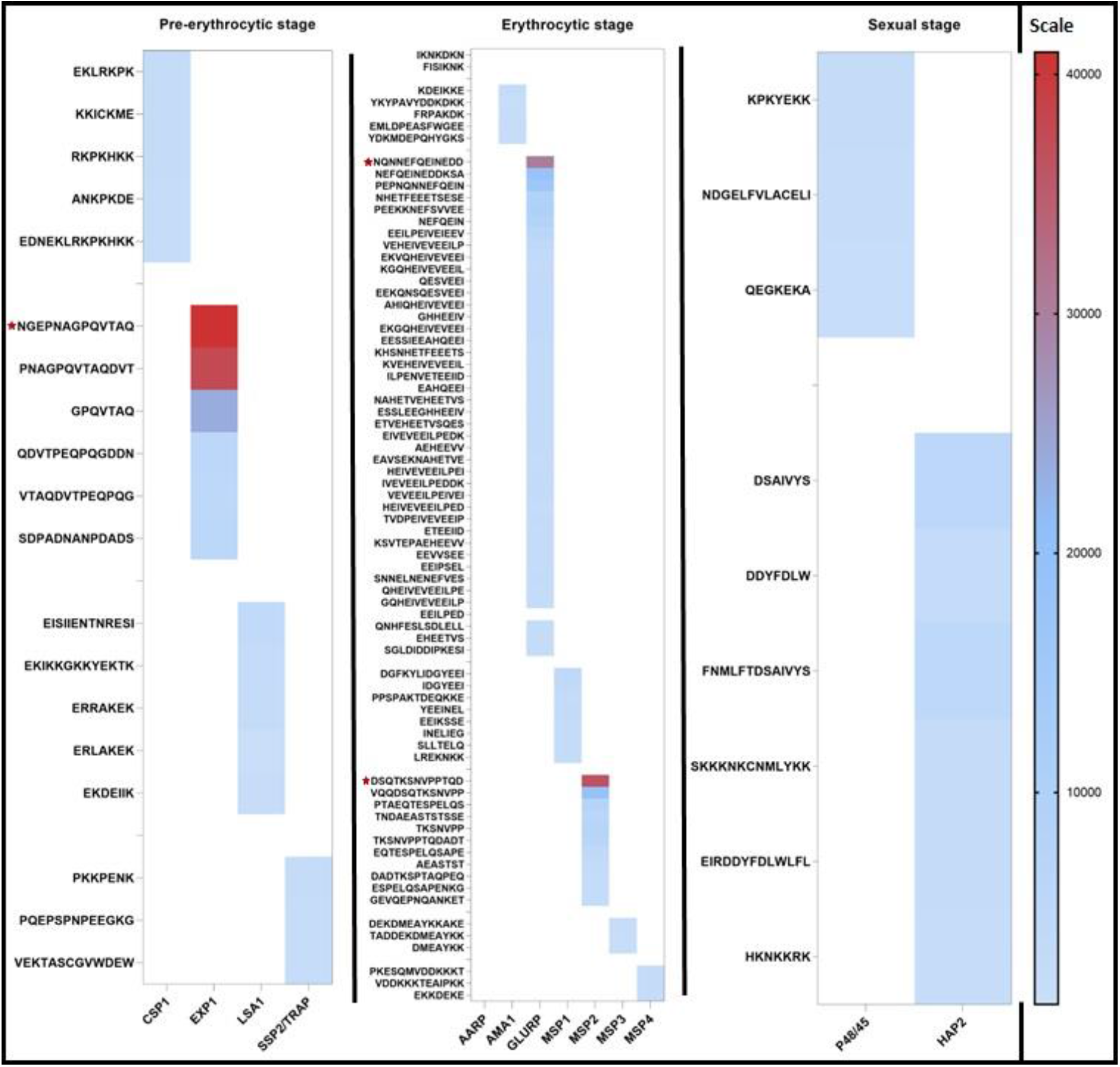
Heat map generated using foreground mean of fluorescence intensities of the representative highest scoring peptides from 13 proteins from multiple stages of the *P. falciparum*. * indicates the three peptides which have shown the highest score amongst all the peptides of the microarray slide.

**Table 2:** List of cyclic constrained and linear peptides chosen for further *in vitro* validation studies.

| S. No. | Protein name | Peptide sequences | Cyclic constrained (C) | Linear (L) |
| --- | --- | --- | --- | --- |
| 1. | EXP1 | NGEPNAGPQVTAQ | C6 | G11 |
| 2. | GLURP | NQNNEFQEINEDD | B7 | NQN |
| 3. | MSP2 | DSQKSNVPPTQD | A8 | DSQ |

#### Conservancy analysis of the shortlisted highly reactive peptides

The conservancy analysis of the shortlisted highly reactive peptides was performed to analyze the presence of these epitopes across different strains of *P. falciparum* malaria parasite. To perform the conservancy analysis, the full-length amino acid sequences of the source proteins (EXP1, MSP2 and GLURP) from different infectious strains of *P. falciparum* were retrieved from NCBI protein database. The multiple sequence alignment (MSA) analysis of the selected epitopes was performed by Clustal Omega on EMBL-EB (https://www.ebi.ac.uk/Tools/msa/clustalo/). The MSA of the selected epitopes show highly conserved nature of the epitopes amongst different strains of *P. falciparum* (Figure 4). Hence the selected epitopes have the potential to be developed as effective and highly specific diagnostic tools.

**Figure 4:**
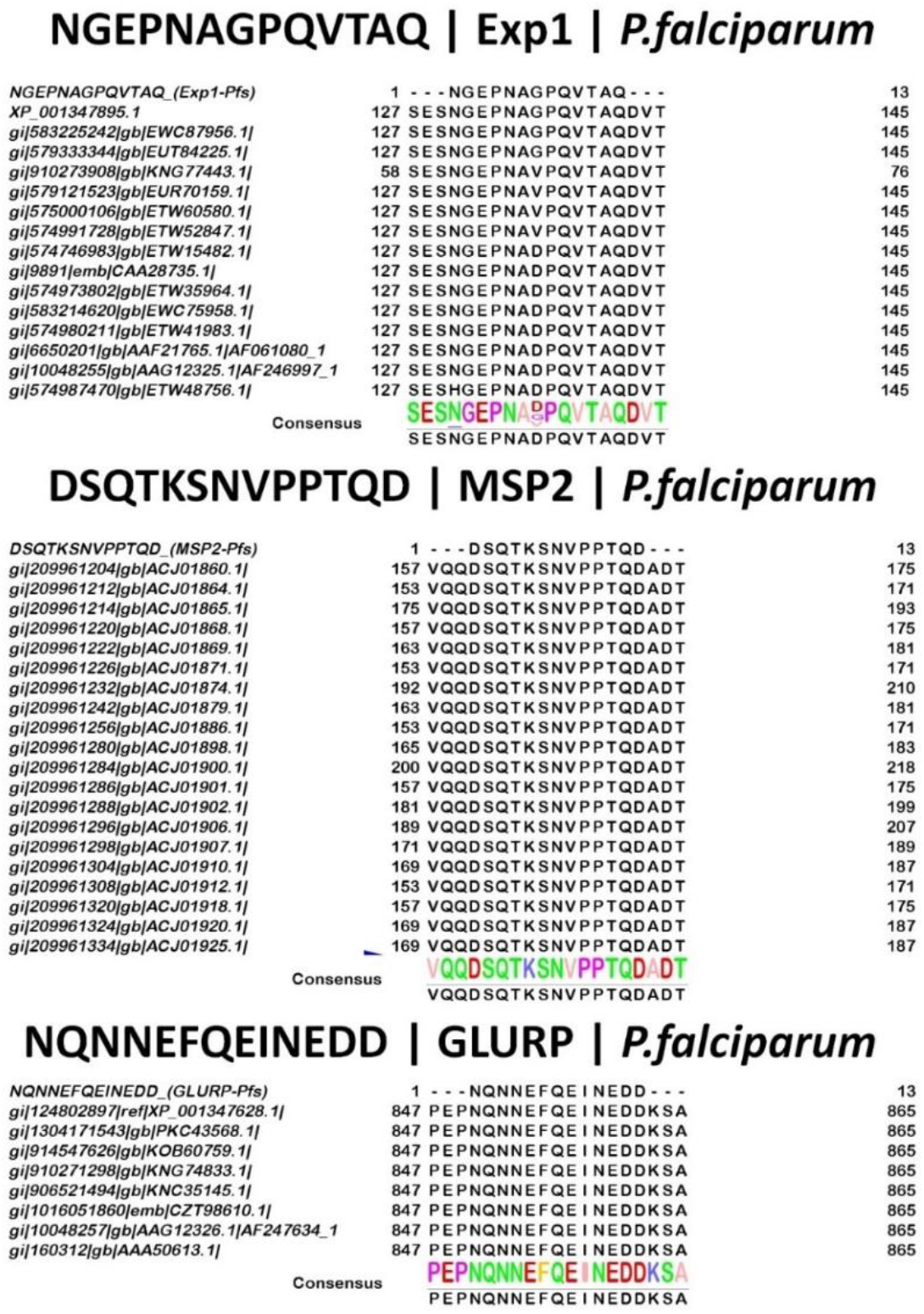
Conservancy analysis of the shortlisted epitopes. The peptides with the highest reactivity were compared by multiple sequence alignment with the source proteins (EXP1, MSP2 & GLURP). The analysis indicates the highly conserved nature of the chosen epitopes. The detailed alignment result of MSP2 could be retrieved from the share [link](https://drive.google.com/file/d/1FLuFS46Xg3GnAjuTHy9-tK2EuwOvTx2x/view?usp=sharing).

### Validation of immunoreactive peptide epitopes

#### Dot blot immunoassay of cyclic constrained and linear epitopes

For preliminary analysis, we included the cyclic constrained and linear epitopes for reactivity against *P. falciparum* infected serum samples. Recombinant glutamate rich protein (GLURP) & *P. falciparum* culture lysate were used as the positive controls. Casein and *P. berghei* culture lysate was used as negative controls. The immunoblots on the PVDF membrane showed visible brown precipitation indicating significant binding and complex formation by the chosen peptides and antibodies in the sera. The qualitative dot blot assay demonstrates that the cyclic constrained peptides from GLURP (B7), EXP1 (C6), MSP2 (A8) generated visible brown precipitate with the DAB substrate indicating their reactivity **(Figure 5)**. Interestingly, two linear peptides GLURP (NQN), P48/45 (KPK) also generated a strong visible brown signal with the substrate. In comparison, the third peptide MSP2 (DSQ) generated a comparatively lighter precipitate in the dot blot assay **(Figure 5)**. Recombinant GLURP used here as a positive control in different concentrations produced visibly strong signals in the dot blot assay. The crude lysate of *P. falciparum* served as an additional positive control. Overall, the chosen three peptides from EXP1, GLURP, & MSP2 proteins, in their cyclic constrained conformation, show a comparative visible signal to that of the positive controls. Notably, both linear and cyclic constrained peptides from EXP1 produced a comparative visible signal to the positive control. Hence, the preliminary analysis conclude the immunoreactivityof the chosen peptides against *P. falciparum* infected patient sera.

**Figure 5:**
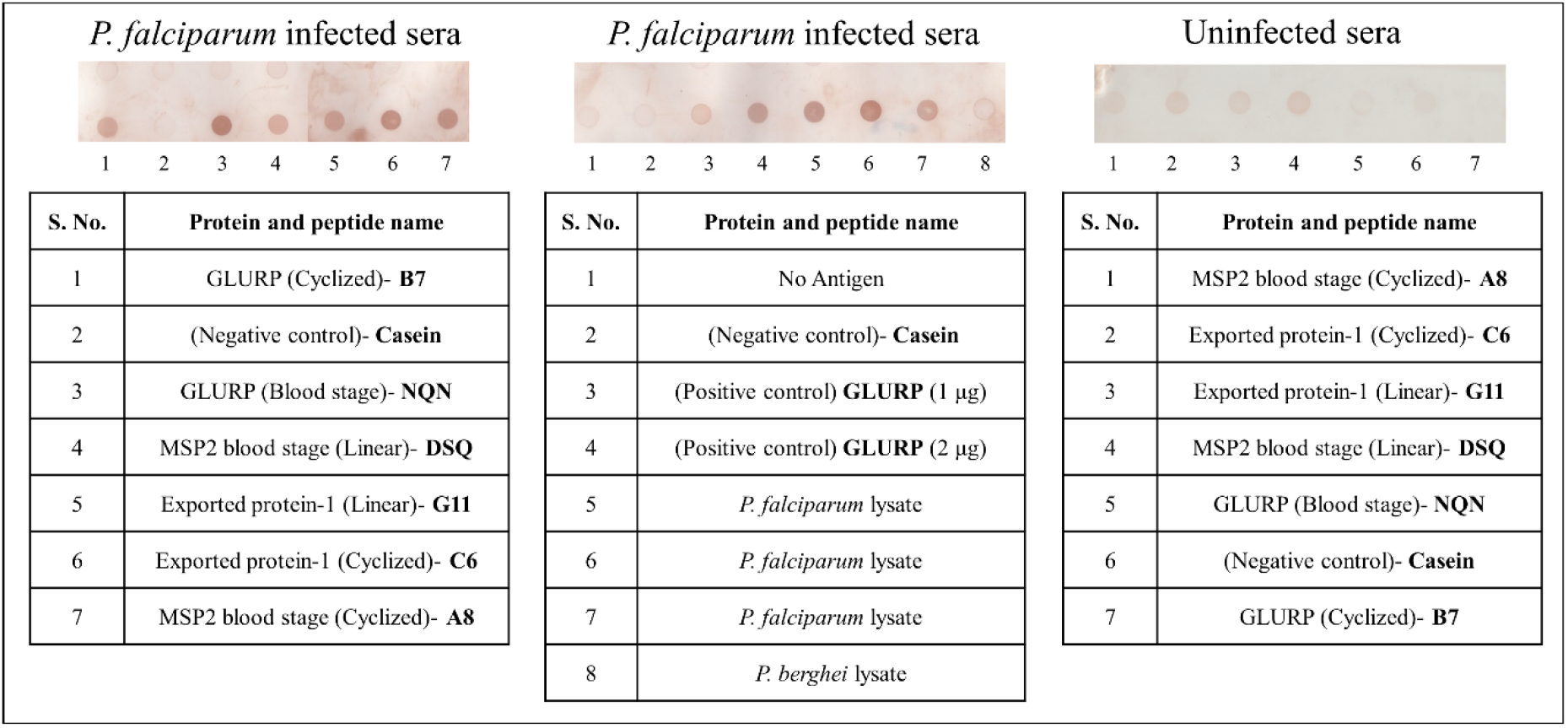
Preliminary dot-blot immunoassay to qualitatively analyse the reactivity of cyclic constrained and linear peptides **(annotations are provided in Table 2)** with *P. falciparum-*infected and uninfected sera.

#### ELISA analysis of cyclic constrained peptides

The cyclic constrained peptides C6 & B7 from EXP1 and GLURP proteins, respectively demonstrated high reactivity with *P. falciparum*-infected patient sera **(Figure 6)**. One of the identified cyclic constrained peptide A8 from MSP2 protein could not be subjected to ELISA, due to very low yield in synthesis. However, there was no reactivity of the peptides towards *P. vivax* infected sera samples, indicating their specificity towards *P. falciparum* **(Figure 6)**. Further, the ELISA experiment was performed with more *P. falciparum-*infected sera samples (n=10) to validate the consistency of the reactive potential of the chosen peptides. The peptides demonstrated significant reactivity towards antibodies from all *P. falciparum*-infected sera samples. The peptides belong to two different life-cycle stages of the *P. falciparum* viz. pre-erythrocyte (EXP1) and erythrocyte stage (GLURP) **(Figure7)**. Together, these results conclude that the cyclic constrained peptides C6 and B7, could serve as potential diangnostic targets, subject to large scale validation studies.

**Figure 6:**
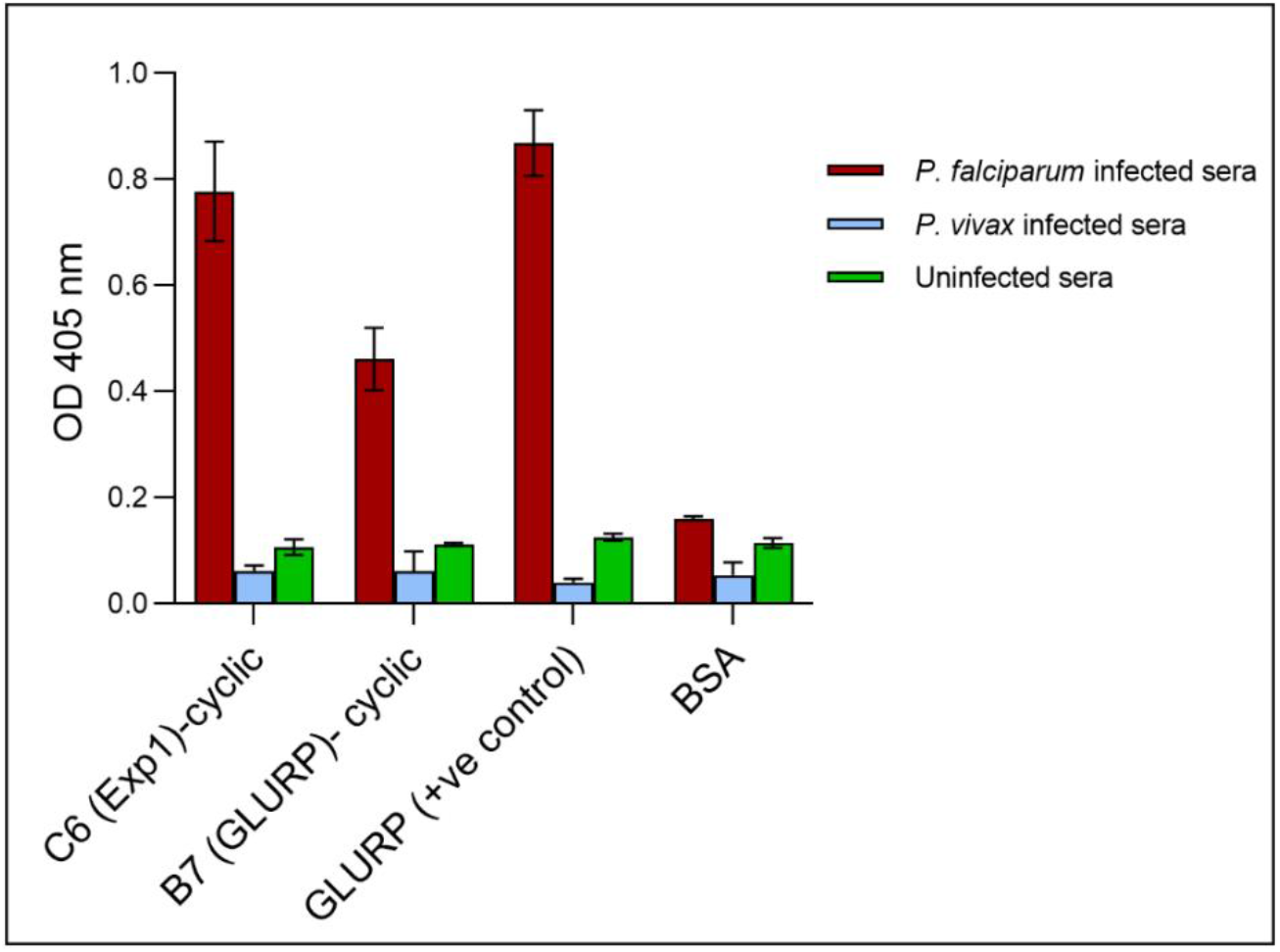
ELISA demonstrating reactivity of cyclic constrained peptides with *P. falciparum, P. vivax* and uninfected sera.

**Figure 7:**
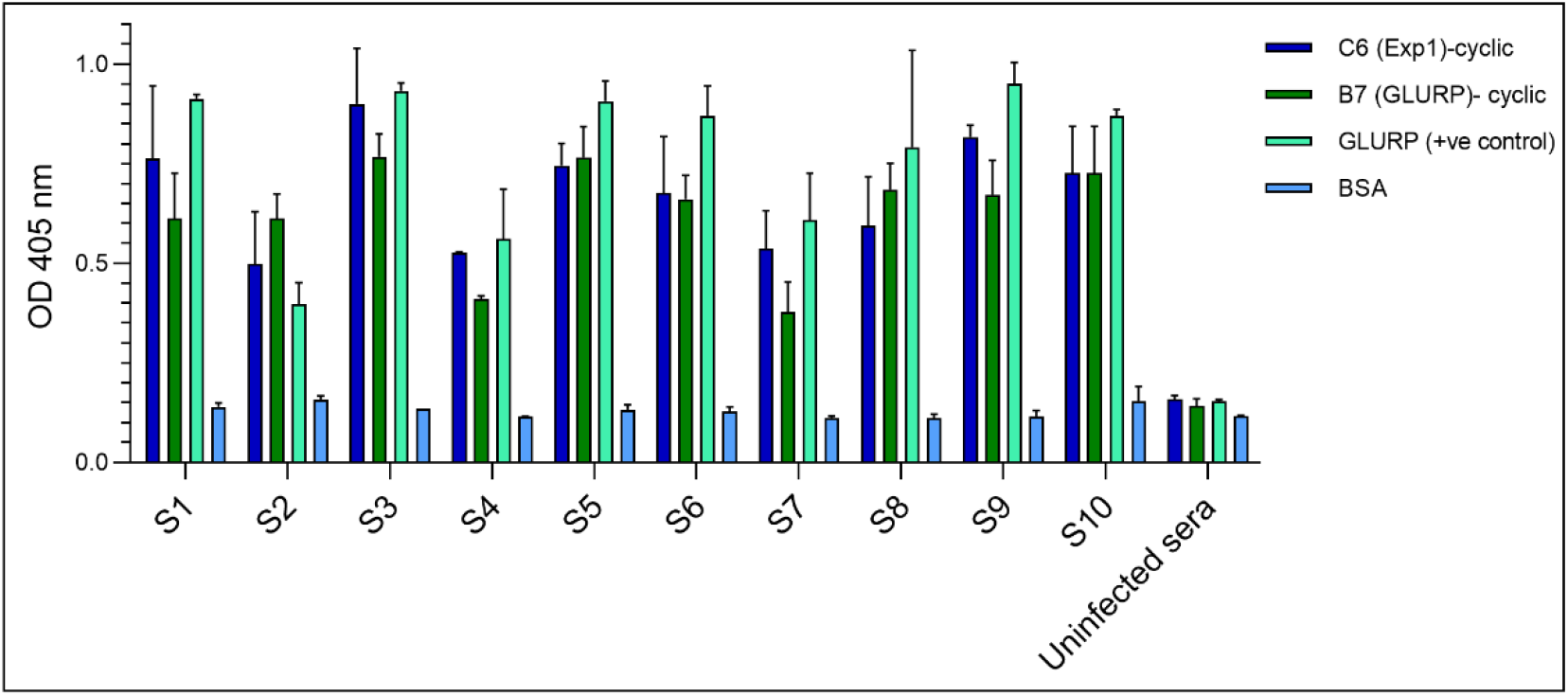
ELISA of cyclic constrained peptides with *P. falciparum* infected sera and BSA as the negative control.

#### Comparison of cyclic constrained peptides vs linear

Notably, the study particularly focused on cyclic constrained peptides using peptide microarray immunoassays. Therefore, it is plausible to assess the comparative reactivity of cyclic constrained and linear peptides. Comparative ELISA between respective cyclic constrained and linear peptides demonstrated that cyclic constrained peptides have significantly high reactivity towards antibodies from *P. falciparum*-infected sera **(Figure 8)**. The better reactivity of the cyclic constrained peptides might be attributed to the 3D mimicry of the epitopes recognized by the antibodies.

**Figure 8:**
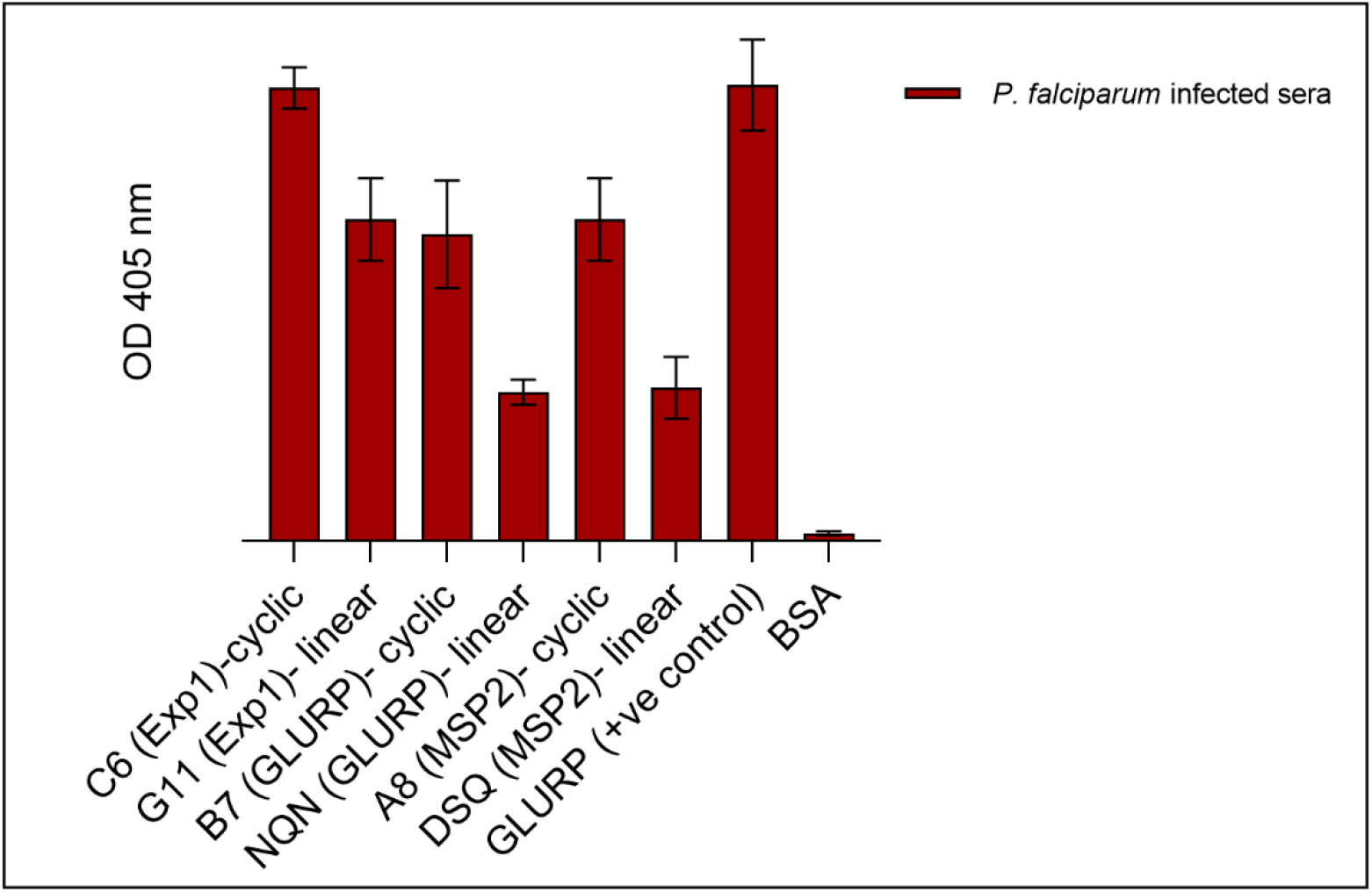
ELISA analysis to compare the reactivity of cyclic constrained and linear peptides with the *P. falciparum*-infected sera.

#### Protein homology model-based analysis of cyclic constrained and linear epitopes

The reason behind the linear peptide C6 (EXP1) being well recognized by the serum antibodies could be suggested by the structural location of the epitope in the protein. Hence, homology models were generated for all three proteins by using I-TASSER web server. The exposure of the selected peptide epitopes in their 3D conformations EXP1, MSP2 and GLURP proteins are shown in **(Figure 9)**. The C6 peptide epitope of the EXP1 is part of a flexible and distant loop exposed towards the surface. Hence, due to its surface exposure, it is easily recognizable by the antibodies. Therefore, the C6 peptide epitope is well recognized in its cyclic constrained and linear conformations. The cyclic constrained peptide epitope B**7** (GLURP) forms an intrinsic loop within the tertiary structure of GLURP, mimicking the natural epitope **(Figure 9)**. Similarly, the location of the peptide epitope A8 (MSP2) in a conformationally structured manner could explain the significant recognition by the antibodies. Thus, the homology models based structural location and conformational analysis of the three peptides corroborates with the findings of the dot blot and ELISA.

**Figure 9.**
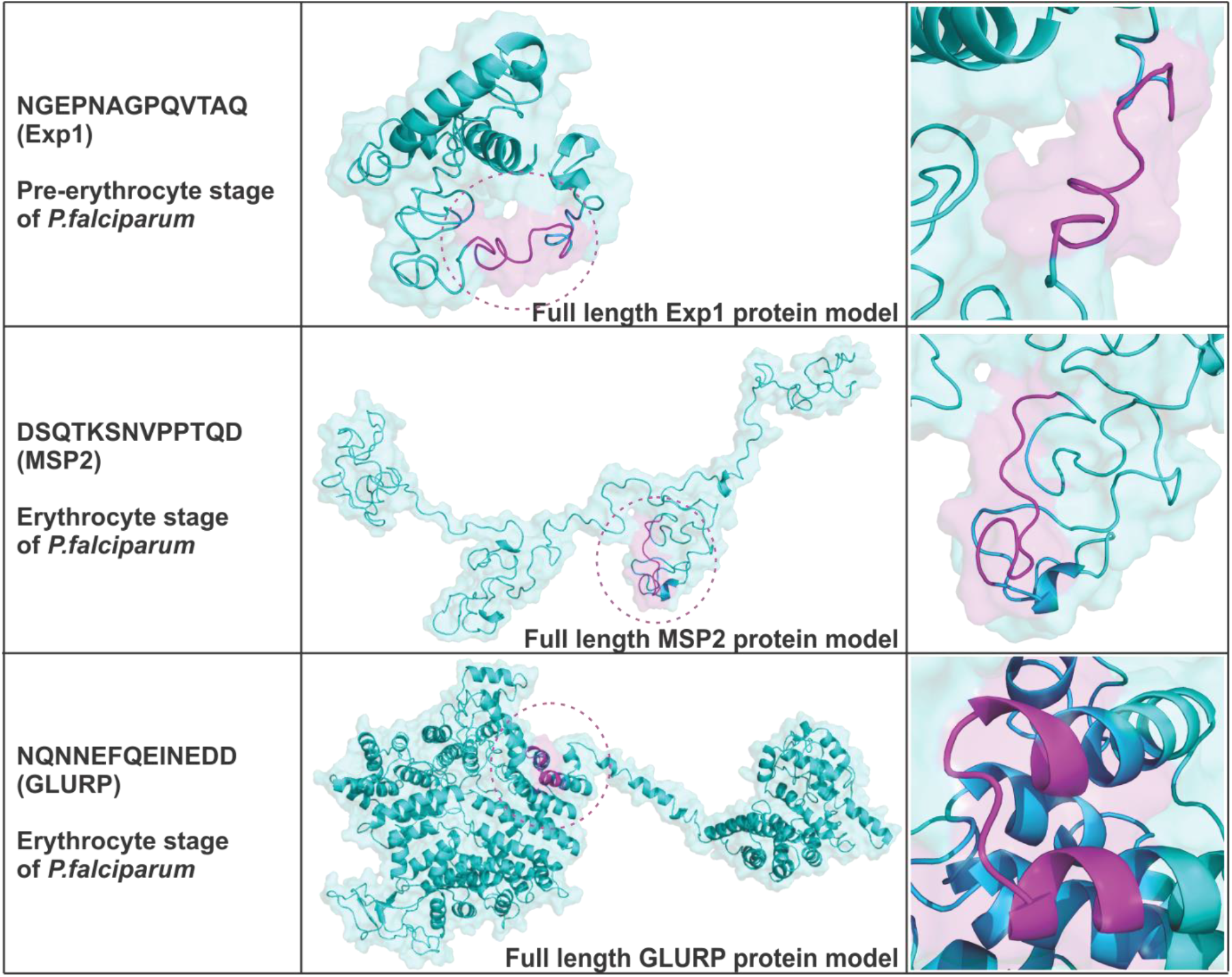
*In-silico* analysis of the chosen peptides in their full-length proteins tertiary structure models. Protein is shown in cyan and the epitope is shown in magenta.

## Discussion

The present study has utilized a scientifically well-established method of peptide epitope mapping using peptide microarray, with immobilized cyclic constrained peptides. The top-scoring three peptides viz. C6 (EXP1), B7 (GLURP) and A8 (MSP2) were identified and further chosen for validation studies. All three selected peptides showed significant visible immunoprecipitation in a dot-blot assay using *P. falciparum-*infected patients sera. The peptides in cyclic constrained conformation have shown substantial visual precipitation indicating favourable complex formation with antibodies of the patient sera. Notably, peptide C6 (EXP1) has shown strong precipitation in its linear conformation as well. Using ELISA, the peptides in cyclic constrained conformation have shown significantly stronger absorbance, indicating that the serum antibodies have a greater propensity to bind to conformational epitopes. Amongst all three shortlisted epitopes, the cyclic C6 (EXP1) peptide has shown the greatest potential for complex formation with antibodies. Notably, the linear peptide G11 (EXP1) also demonstrated a significant response towards antibodies as compared to the other two linear peptides, DSQ (MSP2), and NQN (GLURP). The peptides A8 (MSP2) and B7 (GLURP) also demonstrated significant absorbance in their cyclic constrained conformations, indicating a high propensity of complex formation with the serum antibodies. The linear conformation of the DSQ (MSP2), and NQN (GLURP) peptides demonstrated a relatively weaker absorbance as compared to cyclic constrained conformational peptides. Further, the cyclic constrained peptides C6 (EXP1) and B7 (GLURP) have shown a consistent reactive potential as observed by ELISA validation against ten serum samples of *P. falciparum* infection. Hence, the identified peptides from the present study, may be explored as potential diagnostic targets for *P. falciparum* malaria.

### Conclusion

The present study identified three cyclic constrained peptides viz. C6 (EXP1), B7 (GLURP) and A8 (MSP2), found to be highly immunoreactive to the *P. falciparum* infected sera. The cyclic constrained peptides C6 (EXP1) and B7 (GLURP) were further demonstrated to be specific to *P. falciparum* only. These two peptides were also recognized well by ten field collected samples, indicating their diagnostic potential for *P. falciparum* malaria. The identified peptides C6 and B7 belonged to different stages of the *P. falciparum* life-cycle.Therefore, it would be interesting to assess the multi-stage diagnosis of *P. falciparum* in large scale studies.

**# The results of the present study have been submitted as a patent application at ICMR via application no. 202011015006**.

## Acknowledgment

The authors thank National Research Foundation, South Korea for the funding support. The authors thank PEPerPRINT for the technical support in conducting the peptide microarray study.

## Author contributions

Conceived and designed the experiments: KV, SS and KCP Data generation and analysis: KV, SS, V, SS, NB, AKA, RD Data interpretation, manuscript writing and review: KV, SS, TSK, BKN, HJS, KCP.

## Bibliography

1. World malaria report 2020. https://www.who.int/publications/i/item/9789240015791.

2. Laurent, A. et al. Performance of HRP-2 based rapid diagnostic test for malaria and its variation with age in an area of intense malaria transmission in southern tanzania. Malar. J. 2010 91 9, 1–9 (2010).

3. Makler, M. T., Piper, R. C. & Milhous, W. K. Lactate Dehydrogenase and the Diagnosis of Malaria. Parasitol. Today 14, 376–377 (1998).

4. Bharti, P. K. et al. Prevalence of pfhrp2 and/or pfhrp3 gene deletion in plasmodium falciparum population in eight highly endemic states in India. PLoS One 11, 1–16 (2016).

5. Wangdi, K. et al. Malaria elimination in India and regional implications. Lancet Infect. Dis. 16, e214–e224 (2016).

6. Bartholdson, S. J., Crosnier, C., Bustamante, L. Y., Rayner, J. C. & Wright, G. J. Identifying novel Plasmodium falciparum erythrocyte invasion receptors using systematic extracellular protein interaction screens. Cell. Microbiol. 15, 1304–1312 (2013).

7. Harris, P. K. et al. Molecular identification of a malaria merozoite surface sheddase. PLoS Pathog. 1, 0241–0251 (2005).

8. Kadekoppala, M. & Holder, A. A. Merozoite surface proteins of the malaria parasite: The MSP1 complex and the MSP7 family. International Journal for Parasitology vol. 40 1155–1161 (2010).

9. Boyle, M. J. et al. Sequential processing of merozoite surface proteins during and after erythrocyte invasion by Plasmodium falciparum. Infect. Immun. 82, 924–936 (2014).

10. Baum, J., Maier, A. G., Good, R. T., Simpson, K. M. & Cowman, A. F. Invasion by P. falciparum merozoites suggests a hierarchy of molecular interactions. PLoS Pathog. 1, 0299–0309 (2005).

11. Cowman, A. F., Berry, D. & Baum, J. The cellular and molecular basis for malaria parasite invasion of the human red blood cell. J. Cell Biol. 198, 961–971 (2012).

12. Tan, J. et al. A public antibody lineage that potently inhibits malaria infection through dual binding to the circumsporozoite protein. Nat. Med. 24, 401–407 (2018).

13. Nixon, C. E. et al. Identification of protective B-cell epitopes within the novel malaria vaccine candidate Plasmodium falciparum schizont egress antigen 1. Clin. Vaccine Immunol. 24, (2017).

14. Jaenisch, T. et al. High-density peptide arrays help to identify linear immunogenic B-cell epitopes in individuals naturally exposed to malaria infection. Mol. Cell. Proteomics 18, 642–656 (2019).

15. J, D. et al. Structural basis for recognition of CD20 by therapeutic antibody Rituximab. J. Biol. Chem. 282, 15073–15080 (2007).

